# Structural Insights into Dopamine Receptor-Ligand Interactions: From Agonists to Antagonists

**DOI:** 10.1101/2023.11.03.565579

**Authors:** Emmanuel Barbosa, Heather Clift, Linda Olson, Lan Zhu, Wei Liu

## Abstract

This study explores the intricacies of dopamine receptor-ligand interactions, focusing on the D1R and D5R subtypes. Using molecular modeling techniques, we investigate the binding of the pan-agonist rotigotine, revealing a universal binding mode at the orthosteric binding pocket (OBP). Additionally, we analyze the stability of antagonist-receptor complexes with SKF83566 and SCH23390. By examining the impact of specific mutations on ligand-receptor interactions through computational simulations and thermostability assays, we gain insights into binding stability. Our research also delves into the structural and energetic aspects of antagonist binding to D1R and D5R in their inactive states. These findings enhance our understanding of dopamine receptor pharmacology and hold promise for drug development in central nervous system disorders, opening doors to future research and innovation in this field.

## INTRODUCTION

Dopamine and the dopamine receptor system play critical roles in motor functions, cognition, and addiction [1,2]. Dysfunction of the dopaminergic system has been linked to many central nervous system diseases, including Parkinson’s disease, schizophrenia, and attention deficit hyperactivity disorder (ADHD) [3–6]. Dopaminergic functions are mediated by a family of five G protein-coupled receptors (GPCRs), which are divided into two groups: the D1-like and the D2-like receptors [7,8]. The D1-like group includes D1R and D5R, whereas the D2-like group includes D2R, D3R, and D4R. D1-like receptors are coupled to stimulatory G proteins (G_s_) and linked to the activation of adenylate cyclase. The D2-like receptors are coupled to inhibitory subtypes of G proteins (G_i_ and G_o_) and linked to the inhibition of adenylate cyclase [9].

Recently, Xu et. al [10] published a comprehensive examination of the structures for all known human dopamine receptors. This manuscript was the first to compare the active-state structures of all dopamine receptor complexes bound to the same ligand, rotigotine. Analysis of the structures reveal a universal binding mode of the pan-agonist rotigotine in all five dopamine receptors and the specific intermolecular interactions that define the recognition of rotigotine by each of them. Structural and sequence comparisons indicate that the conserved orthosteric binding pocket (OBP) is the basis for promiscuous binding of rotigotine in each of these receptors.

In this paper, we explore these OBP interactions with the agonist rotigotine, employing molecular modelling techniques to detail the atomic-level consequences of single point on ligand binding. Additionally, a study with inactive structures binding antagonists is performed.

Binding of a ligand to a GPCR can affect the protein’s dynamics in several ways. Generally, the largest effect is simply to alter the fraction of time the GPCR spends in each of its conformational states. For example, binding of certain ligands may increase the population of active conformational states or increase/decrease the rate at which the receptor transitions among different conformational states [11]. Determining these atomic fluctuations is very computationally intensive, but atomic-level molecular dynamics (MD) simulations have become a powerful complement to traditional structural methods such as crystallography and spectroscopy in recent years. Increased computer power, improved simulation algorithms, and refined potential energy functions that better represent the underlying physics have allowed for this [12,13]. Starting from a static experimental structure, these simulations predict the motion of every atom in a receptor, as well as the motion of every atom in the molecules with which the receptor interacts [14].

Currently, there are only active-state structures for D1R and D5R (agonist-bound), so little is known about the inactive structures (antagonist-bound). The basic definition of agonist and antagonist in GPCRs is as follows: Agonists are defined as the ligands that activate intracellular signaling and evoke cellular responses. They bind specific amino acids to activate the GPCR. Antagonists inhibit agonist-stimulated responses, usually by competitively binding to orthosteric or allosteric sites, and are often related with inverse agonist activity [15]. However, as with most things in nature, agonist and antagonist binding and activation/inactivation is more complex and predominantly relies on the specific cellular situation. For example, an “antagonist” could actually be acting as an inverse agonist, partial agonist, or a neutral antagonist, and may not even bind directly to the receptor, blocking activation of the GPCR by disrupting its interaction with the G protein [16]. It is therefore important to obtain specific receptor-ligand interaction data, including computational models, assay results, and protein structures.

Focusing on D1R and D5R structures, we have selected specific OBP alanine mutants from the published rotigotine paper [10] and compared our computational results with their cAMP assays. Based on these results, we employed molecular modelling techniques to docking and predict antagonist binding of SKF83566 and SCH23390 to both D1R and D5R, followed by the production of experimental investigation.

## MATERIALS AND METHODS

### Molecular Simulation

To perform the *in silico* simulations of the Dopamine receptors in the active state, we use the atomic coordinate data of the wild-type human Dopamine receptors D1 and D5 complexed with the agonist rotigotine obtained from PDB ID 8IRR and 8IRV, respectively[10]. Due to the lack of active-state solved structures for D1 and D5 receptors, [AlphaFold Protein Structure Database](https://alphafold.ebi.ac.uk/) [17] was used to generate the corresponding inactive models for D1 and D5; The N-terminal and C-terminal regions of the inactive state with a low level of confidence were removed, and the inactive structure models assume the same amino acid sequence of the active structures solved by Cryo-EM method. The molecular docking of the antagonists SCH23390 and SKF83566 within the inactive models’ binding pocket was performed using Autodock Vina Software [18]. After obtaining the protein–ligand complexes, the protonation state study of the ligands and protein amino acids at pH 7.4 were performed using the Marvin Sketch code, version 18.24 (*ChemAxon, https://www.chemaxon.com*), and the [19] PROPKA 3.1 package, [20] respectively. The topology and coordinates files of the Dopamine receptors embedded in a lipid bilayer were generated using with the Charmm-GUI web server [21]. The main components of a complex neuronal/synaptic membrane were simulated using a molar ratio of 5:3:2, for cholesterol, 1-Palmitoyl-2-oleoyl-phosphatidylcholine (POPC), and 1-palmitoyl-2-oleoylphosphatidylethanolamine (POPE) [22]. The atomic partial charges of the ligands were computed by using the ambertolls (v.2021)/antechamber package [23]. The GROMACS software (v.2020.5) [24] was used to run all the molecular dynamics simulations under the Charmm36m force-field along 40 ns. The temperature of 310 K and *v*-rescale thermostat algorithm are used during the production run.

The Molecular Dynamics-protein-ligand Interaction FingerPrints (MD-IFP) [25] method was used to characterize the intermolecular interactions between receptor and ligand atoms computed using the last 1000 frames from 30-40 ns, saved at intervals of 10 ps for each trajectory simulation, where the prefixes AR, HY, IP, HD, and HA correspond to Hydrophobic, Ionic Positive, Hydrogen-Bond Donnor, and Hydrogen-Bond Acceptor interaction, respectively.

The Quantum/Molecular Mechanics Generalized Born Surface (QM/MMGBSA) approach [23] was employed to obtain the protein-ligand binding free energy values (in Kcal/mol). The semiempirical quantum chemical PM6 theory level with additional terms for dispersion and hydrogen bonding was applied for the residues within 5 Å of the ligand, and the classical amber force field ff19sb for the remaining protein amino acids. The binding affinity between the dopamine receptor and ligands was measured using 100 snapshots from 30ns to 40ns saved at intervals of 100 ps of each trajectory.

### D1R Expression and Purification

Human D1R wild-type was synthesized and cloned into the expression vector pFastBac1 containing an expression cassette with an HA signal sequence followed by a FLAG tag at the N-terminus and a PreScission protease site followed by a 10xHis tag at the C-terminus. The components of the expression cassette were introduced using standard PCR-based site-directed mutagenesis. D1R mutants were modified by introducing every single mutation using standard QuickChange PCR and then sub-cloned into the pFastBac1 vector. High-titer recombinant baculovirus was obtained using the Bac-to-Bac Baculovirus Expression System (Invitrogen), and used to infect sf9 insect cells to produce D1R proteins as described previously [26]. Cells were harvested by centrifugation at 48 hours post infection and stored at -80 °C until use.

D1R wild-type and mutants were purified in complexes with rotigotine [26]. Thermostabilities were examined in a detergent solubilized state using a thiol-specific fluorochrome N-[4-(7-diethylamino-4-methyl-3-coumarinyl) phenyl]maleimide (CPM) assay in which fluorescence of a maleimide on binding to free CYS sulfhydryls is monitored as temperature is raised and the receptor unfolds [27].

## RESULTS

Xu et al. recently published a comprehensive study of the dopamine family of receptors bound to rotigotine. They reported that even though D1 and D5 had essentially identical backbone conformations (root-mean-square deviation 0.48 Å) and the orthosteric and extended binding pockets (EBP) were highly conserved. Furthermore, they further characterized the effects of alanine substitutions on the potency of D1R and D5R towards rotigotine. Based on their findings, molecular modeling studies were initiated on residues around rotigotine to investigate the binding mechanisms between D1R (D5R) and rotigotine (Fig. 1). A general view and overlayed structures depicting the identified amino acids residues are also presented (Fig. 1AB). Results regarding the type and frequency of the intermolecular interactions are further annotated in Fig. 1C, where we also reference the Ballesteros-Weinstein sequence to provide a comprehensive view of the residues involved in the ligand binding and recognition process. We noted that in D1R, the amino acids W99^D1R^ and V317^D1R^ presented more frequent hydrophobic interactions compared to their counterparts in D5R and the residues, L190^D1R^, F288^D1R^ and W321^D1R^, showed a minor difference in occurrence. While in D5R, F341^D5R^ exhibited a higher hydrophobic interaction frequency than its counterpart in D1R.

**Figure 1.**
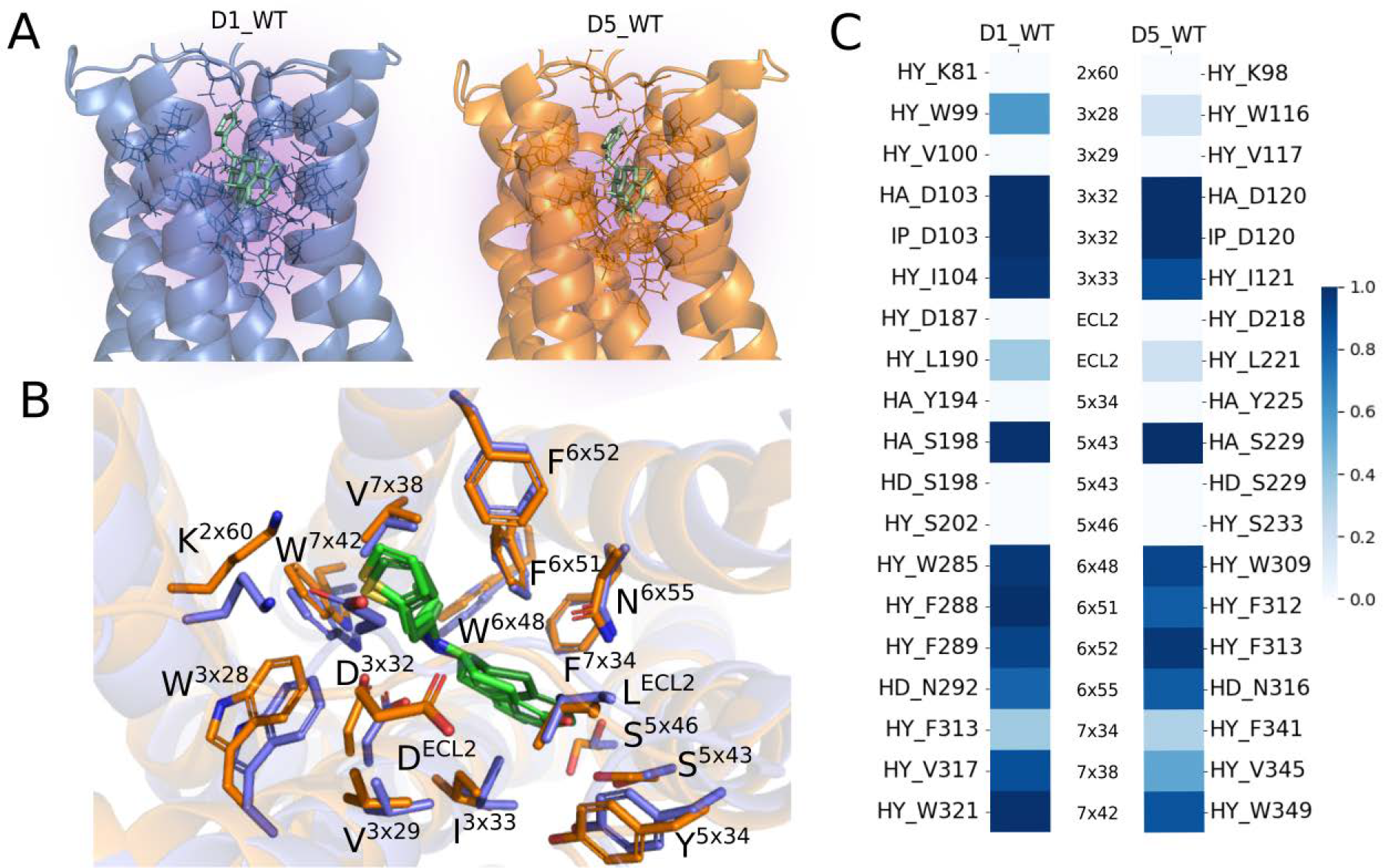
Comparative overview of the wild-type dopamine D1 receptor family: A) Schematic illustration of the Wild-Type (WT) Dopamine receptors D1 (blue) and D5 (orange) binding to Rotigotine (green) from PDB IDs 8IRR and 8IRV, consecutively; The surrounding residues from Rotigotine are outlined. B) Top-view of rotigotine binding pocket of D1R and D5R, highlighted amino acids consistent with the comparison in panel C. C) Graphic panel showing the amino acids interacting with rotigotine, the abbreviation of the intermolecular interaction type, and its occurrence; The abbreviations AR, HY, IP, HD, HA and refer to Aromatic, Hydrophobic, Ionic Positive, Hydrogen-Bond Donnor and Hydrogen-Bond Acceptor interaction, respectively; The Ballesteros-Weinstein sequence is present connecting the correspondent residues.

To further investigate the robustness of our method, six residues were selected for *in silico* mutation in D1R and D5R based on the changes in the potency to stimulation with rotigotine results published by Xu *et al*. (Figs. 2 and 3) [10]. First D103^D1R^/D120^D5R^, located in helix 3 within the OBP, was reported to have substantial decreases in potency to rotigotine when mutated to alanine in both receptors. Also selected S202^D1R^ /S233^D5R^ located in helix 5 and previously identified to be involved in both D1R and D5Rs’ binding to dopamine but not highly relevant to rotigotine, showing either a non-significant or slight increase in potency in this assay and K81^D1R^/K98^D5R^ previously identified in binding tavapadant and PW0464 and showing no changes in stimulation with rotigotine. Two residues with opposing sensitivity to rotigotine when mutated to alanine (F313^D1R^ /F341^D5R^ and V317^D1R^ /V349^D5R^) were also selected. Finally, one residue in the EBP (L190^D1R^ /L221^D5R^) demonstrating a decrease in potency was also mutated to alanine for further investigation.

**Figure 2.**
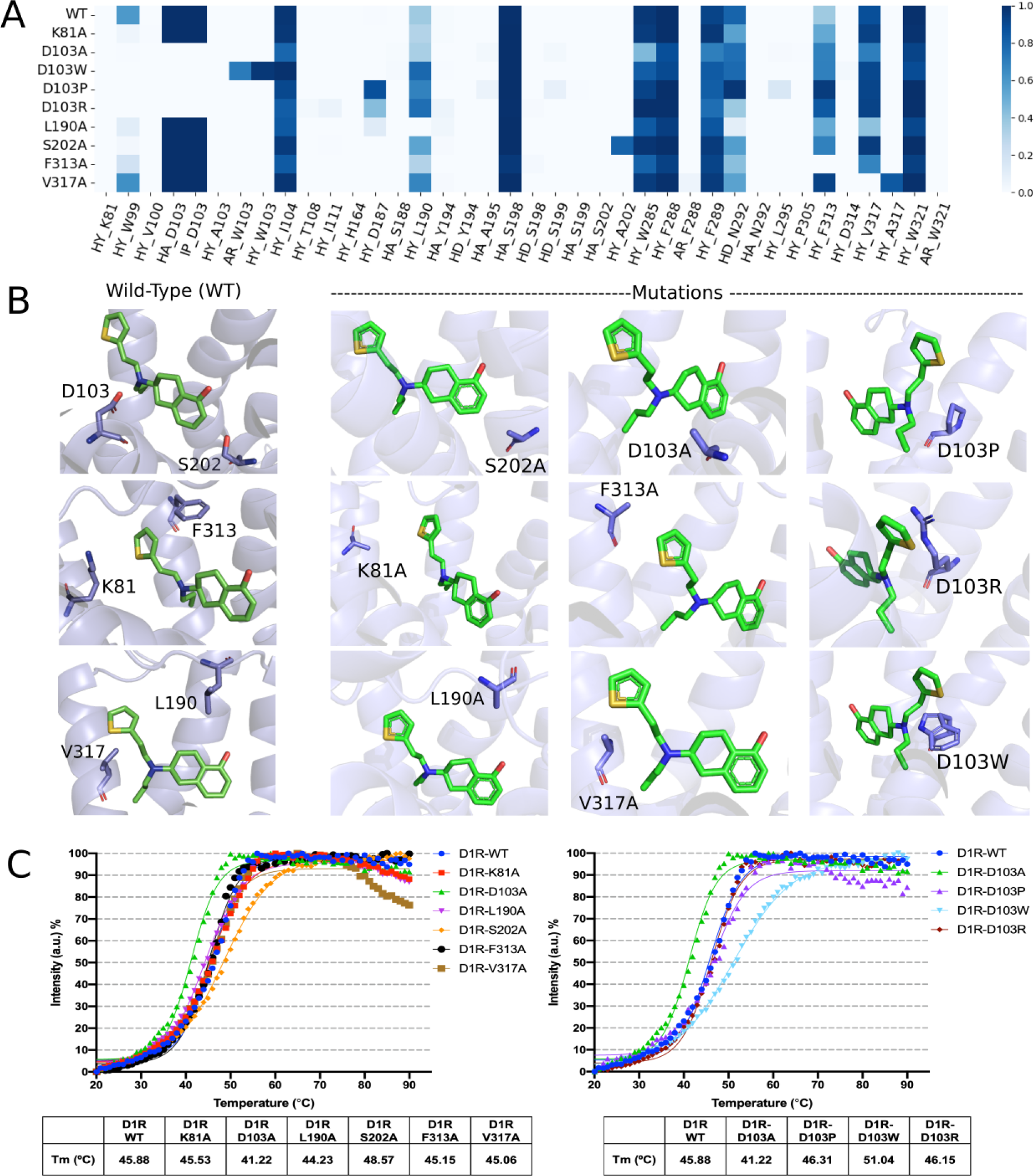
Visualization of the Wild-Type (WT) Dopamine D1 Receptor Compared to Its Mutants: A) A heatmap displays the residues interacting with rotigotine, the type of intermolecular interaction (abbreviated), and their frequency for both the WT D1 receptor and its mutants: K81A^D1R^, D103A^D1R^, D103W^D1R^, D103P^D1R^, D103R^D1R^, L190A^D1R^, S202A^D1R^, F313A^D1R^, and V317A^D1R^. B) Depiction of the WT residues (right side) and the impact of mutations (left side) on the ligand-binding pocket of D1R. C) Thermostabilities of D1R WT and mutants proteins bound with rotigotine. Boltzmann sigmoidal statistics used to create nonlinear curve fitting from normalized intensity, the calculated Tm (°C) shown in the table.

**Figure 3.**
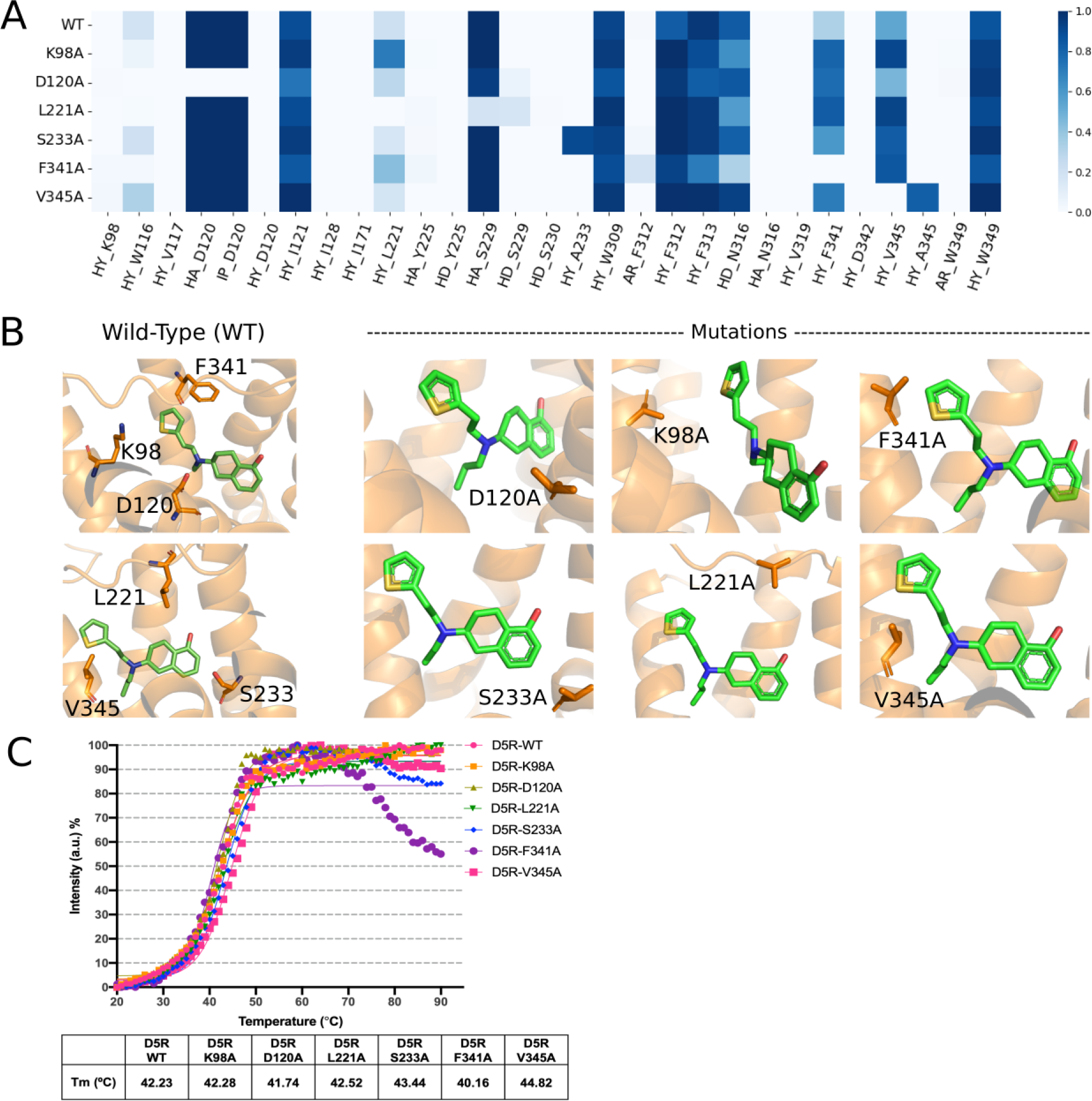
Illustration of the Wild-Type (WT) Dopamine D5 Receptor Compared to Its Mutant variations: A) A heatmap showcases the amino acids interacting with rotigotine, the type of intermolecular interaction (abbreviated), and occurrence for both the WT D5 receptor and the mutations: K98A^D5R^, D120A^D5R^, L221A^D5R^, S233A^D5R^, F341A^D5R^, and V345A^D5R^. B) Effect of mutations on the ligand-binding pocket of D5R. C) Thermostabilities of D5R WT and mutants proteins bound with rotigotine. Boltzmann sigmoidal statistics used to create nonlinear curve fitting from normalized intensity, the calculated Tm (°C) shown in the table.

A heatmap panel was used to present the interacting residues, types of intermolecular interactions, and their frequencies for both the WT D1R and the (Fig. 2A). Mutants K81A^D1R^, F313A^D1R^, and V317A^D1R^ displayed little to no significant differences in the interaction heatmap, with rotigotine implying the shape of the OBP and position of the ligand are maintained even with the mutation. D103A^D1R^ and L190A^D1R^ mutants exhibited a notable inability to interact with rotigotine in the pocket. Remarkably, rotigotine demonstrated a high frequency of interactions with D1R at the mutation S202A^D1R^ positioned on the transmembrane helix 5 (TM5), the non-polar residue (alanine) showed able to provide a high frequency of hydrophobic interaction. Additionally, interactions at L190^D1R^ and F313^D1R^ were enhanced via hydrophobic effects. Changes observed from these mutants matched to the ligand efficacy profile from the Gs-mediated cAMP assay results of D1R in previous publication [10]. Extended mutation analysis at D103^D1R^ in our study further characterized its impact on ligand-receptor binding stabilities. Despite the mutation D103W^D1R^, rotigotine could still bind to D1R, primarily via aromatic and hydrophobic effects; furthermore, the interactions were strengthened at residues L190^D1R^, F313^D1R^, and V317^D1R^ via hydrophobic effects. Whereas the interaction patterns of D103^D1R^ mutations to P (Pro) and R (Arg) were differentiated that interactions at other residue bindings were semi-occasional; nevertheless, an increase were noted at D187^D1R^, L190^D1R^, W285^D1R^, and F313^D1R^ via hydrophobic effects. This analysis sheds light on how these mutations impact the ligand-binding pocket of D1R, with each mutation having a unique effect on the receptor’s binding characteristics.

Figure 3 focuses on the dopamine D5 receptor and its mutants. We utilize a heatmap to showcase the interacting residues, types of intermolecular interactions, and their occurrences for both the wild-type D5 receptor and its mutations, including K98A^D5R^, D120A^D5R^, L221A^D5R^, S233A^D5R^, F341A^D5R^, and V345A^D5R^. It revealed various effects of D5R mutants on rotigotine interactions. Mutants D120A^D5R^, K98A^D5R^, and L221A^D5R^ had little to no significant differences in ligand binding. Mutant S233A^D5R^ exhibited enhanced interactions in the same site compared with the wild-type residue and stronger binding with rotigotine, particularly at F312^D5R^ and F341^D5R^. Mutant F341A^D5R^ weakened ligand binding to D5R. Mutant V345A^D5R^ demonstrated notably stronger interactions with rotigotine, with enhanced interactions at F312^D5R^ and F341^D5R^. These findings validated the potency profile [10] and demonstrated different patterns on the pharmacological aspects of rotigotine binding to D5R versus D1R. By comparing the interaction profiles of the mutants with the WT D5 receptor, we gain a comprehensive understanding of the specific effects of these mutations on the receptor’s ligand-binding pocket.

Molecular dynamic simulations provided valuable data on the interaction occurrence frequencies within the ligand-receptor binding pocket. We found that certain mutants exhibited increased interaction occurrence frequencies, indicating enhanced stability in the ligand-receptor binding pocket. Remarkably, this increase in interaction frequency was consistent with higher thermostability observed in the thermostability assays (Fig. 2C, 3C). Conversely, mutants that displayed reduced interaction occurrence frequencies in the binding pocket exhibited decreased thermostability. This observed correspondence between interaction frequencies and thermostability demonstrates that the ligand-receptor binding stability is closely linked to the receptor overall thermostability.

In order to further elucidate the D1 and D5 receptors’ activation process, we studied the inactive state structures of wild-type dopamine receptors D1 and D5 in complex with the antagonists SCH23390 and SKF83566 (Fig. 4). Panel A illustrates the structural representation of SCH23390 (in magenta) bound within the D1 receptor (in blue) and SKF83566 (in cyan) binding to the D5 receptor (in orange). Both antagonists, SCH23390 and SKF83566, are shown interacting with their respective receptors, with amino acids within a 5Å radius of the antagonist compounds depicted. Panel B displays energy profiles detailing the binding interactions between the antagonists and the D1 and D5 receptors. These profiles are generated based on 100 molecular dynamics snapshots using the QM/MM method. These energy profiles provide crucial information about the stability of the antagonist-receptor complexes and the underlying molecular forces governing these interactions. Panel C offers a graphical representation of the amino acids involved in interactions with the antagonists, along with the respective types of intermolecular interactions and their frequency. It is important to mention that the MD-IFP algorithm used in this work identify the hydrogen bonds by considering both the Hydrogen bond Acceptor (HA) and Donnor (HD), which may vary in proportion in the heatmap panel depending on the type of hydrogen bond detected. Thus, some residues can present hydrogen bonds classified as HA and HD or just as one type, HA or HD. The amino acid residues are linked to the Ballesteros-Weinstein sequence, providing a comprehensive view of the specific residues contributing to the binding of SCH23390 and SKF83566 with D1 and D5 receptors. Both antagonists possess a similar molecular structure, which reflects a consistent interaction pattern for both receptors due to their notable homology. The most prominent difference possibly related to the order of affinity of the antagonists is that SCH23390 shows high occurrence of hydrogen bonds in the site S198^D1R^/S229^D5R^ than SKF83566 for both receptors. At the same hand, the non-polar interactions facilitated by nearby residues F288^D1R^/F312^D5R^ and F289^D1R^/F313^D5R^ could co-occur with the previous ones, highlighting the molecular features associated to the binding profiles of each complex. Overall antagonist binding profiles were also validated by ligand-receptor thermostability analysis (Fig. 4D). The data presented in Figure 4 significantly enhances our understanding of the structural and energetic aspects of antagonist binding to dopamine receptors, shedding light on the specific molecular determinants governing these interactions in their inactive states. This knowledge is crucial for the development of novel therapeutics targeting these receptors and may aid in the design of more effective drugs for various central nervous system disorders.

**Figure 4.**
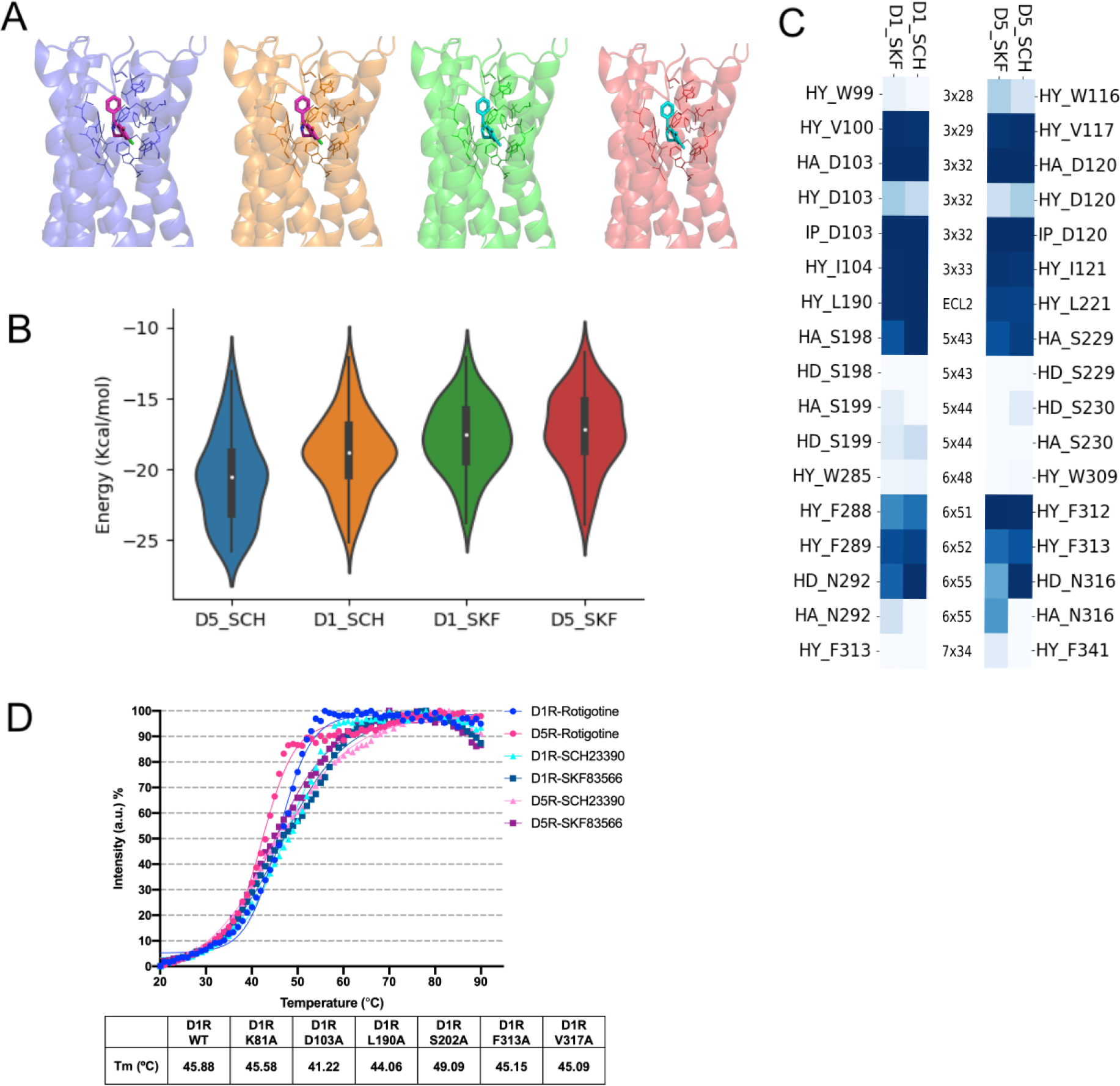
Inactive state structures of Wild-Type Dopamine receptors D1 and D5 binding with SCH23390 and SKF83566 antagonists A) Representation of SCH23390 (magenta) within the D5 (blue) and D1 (orange) receptor and SKF83566 (cyan) binding to D1 (green) and D5 (red) receptor in green and red, respectively; Amino acids within a 5Å radius of the antagonist compounds are depicted. B) Binding energy profiles detailing antagonist’s binding interactions with D1(D5), generated from 100 molecular dynamics snapshots using the QM/MM method. C) Graphic panel depicting the amino acids involved in interactions, the respective intermolecular interaction types, and their frequency. The corresponding residues are also linked to the Ballesteros-Weinstein sequence.” D) Thermostabilities of D1R and D5R WT proteins bound with 3 different ligands. Boltzmann sigmoidal statistics used to create nonlinear curve fitting from normalized intensity, the calculated Tm (°C) shown in the table.

## DISCUSSION

The results presented in this study, along with the insights provided by the figures, offer a comprehensive understanding of the interaction between dopamine receptors and their ligands, including both agonists and antagonists. We have analyzed the structural and energetic aspects of these interactions, shedding light on key molecular determinants and implications for the pharmacology of dopamine receptors. The results presented in Figures 1, 2, 3 and 4 provide critical insights into the interaction between dopamine D1 and D5 receptors and the ligand rotigotine. The comparative analysis of wild-type receptors and mutants demonstrates the influence of mutations on the binding properties of these receptors. Notably, we have observed a universal binding mode of rotigotine in both D1 and D5 receptors, underlining the significance of these findings in the context of dopamine receptor pharmacology.

Our computational analysis showcases the diverse effects that specific mutations have on receptor-ligand interactions. This knowledge is pivotal in advancing our understanding of the precise molecular determinants governing receptor-ligand binding. Additionally, this information contributes to the broader field of GPCR pharmacology, providing a foundation for designing ligands with enhanced selectivity and affinity. Moving forward, we intend to extend our computational models to predict antagonist binding of SKF83566 and SCH23390 to both D1R and D5R. By doing so, we aim to complement our structural insights with predictions of how antagonists may interact with these receptors. These predictions will guide our subsequent efforts to produce experimental structures of the inactive (antagonist-bound) states, further enhancing our understanding of dopamine receptor dynamics.

This study presents a comprehensive investigation into the stability of ligand-receptor binding interactions in a target receptor with various mutants. The study employs a combination of molecular dynamics simulations and thermostability assays to shed light on the impact of receptor mutations on the binding pocket’s stability and interaction frequency.

The molecular dynamics simulations offer valuable insights into the dynamic behavior of the ligand-receptor binding pocket. By quantifying the occurrence frequencies of these interactions, the study demonstrates that specific mutants exhibit increased binding stability when compared to the wild-type receptor. This finding is further reinforced by the analysis of thermostability, which was assessed using a detergent solubilized state and a thiol-specific fluorochrome assay (CPM).

Remarkably, the results of the thermostability assays show a clear correspondence with the molecular dynamics simulations. Mutants that exhibit enhanced interaction occurrence frequencies in the simulations also demonstrate increased thermostability, while those with reduced binding pocket stability exhibit lower thermostability. This consistency between the two analytical approaches suggests a strong correlation between enhanced ligand-receptor binding stability and increased thermostability in mutant receptors.

In summary, this study provides a compelling and robust foundation for understanding the impact of receptor mutations on ligand-receptor binding interactions and how these changes manifest in the thermostability of the receptor. It offers valuable insights into the molecular basis of dopamine receptor-ligand interactions, paving the way for the development of more selective and potent compounds targeting these receptors. These findings contribute to the broader understanding of GPCR pharmacology and hold promise for the development of novel therapeutics for various central nervous system diseases and disorders. The researchers also enhance our understanding of dopamine receptor pharmacology by providing detailed structural, energetic, and mutational insights into the binding of ligands, including both agonists and antagonists. The findings presented here contribute to the broader field of GPCR pharmacology, offering a foundation for the design of ligands with enhanced selectivity and affinity, and providing valuable knowledge for the development of therapeutics for central nervous system diseases and disorders. The combination of computational models, assay results, and experimental studies presented in this work is crucial for deciphering the complex interactions between receptors and ligands and holds great promise for future drug discovery efforts.

## CONCLUSION

Among the aminergic GPCRs, the D1 family of dopamine receptors have ascendent interest due its involvement in both neurological and psychiatric disorders. Therefore, considerable efforts have been made to characterize these receptors and its ligand binding mechanisms to enable the optimization of the D1 family activity, probably leading in improved therapeutic outcomes. However, the current state of treatments based on D1 family receptors demonstrate that there is still a need to unveil the action of agonists and antagonists targeting D1 family receptors, which can enhance its therapeutic potential and minimize adverse effects. In the present study, by taking advantage of recent published structures of the D1 family of dopamine receptors in complex with the agonist rotigotine, and experimental assays reporting the effect of punctual mutation at the orthosteric binding pocket (OBP) of these receptors, we have explored the binding mechanisms of 9 dopamine D1 receptor mutants and 6 dopamine D5 receptor mutants with rotigotine by computational and experimental approaches. First, we studied the OBP from the wild-type receptors interacting with rotigotine to obtain the intermolecular interaction occurrence. Our computational results show that D1R OBP reached higher values of affinity than D5R and the residues W99^D1R^ and V317 ^D1R^ from D1R showed relevant to the most frequent hydrophobic contribution.

In addition, we compared the wild-type receptors with the chimeras to investigate the effects of each mutation into OBP binding rotigotine and the behavior of the intermolecular interactions co-occurring during the molecular dynamic simulation. For both receptors, the mutation S202A^D1R^/S233A^D5R^ showed advantageous by virtue of the elevated occurrence of intermolecular interactions of OBP residues with rotigotine. This enhancement facilitates the improvement of ritigotine-TM5 binding affinity and longer ligand residence time, clarifying the high activity response of this mutation within the D1 family of dopamine receptors.

Finally, an investigation with inactive structures, showed that the antagonists SKF83566 and SCH23390 presented a similar occurrence of chemical interactions and binding energy profiles, with the compound SCH23390 obtaining slight advantageous compared with SKF83566 binding to the D1 family of dopamine receptors.

Our results, therefore, are helpful to elucidate, at the molecular level, the binding features of the agonists and antagonists with respect to the D1 family of dopamine receptors, highlighting the interacting amino acids investigated with the purpose to improve the understanding of the mechanism of GPCRs and facilitate the development of new chemicals targeting dopamine receptors.

Future Research Directions: The results of this research open up avenues for future investigations. The understanding of the role of dopamine receptors in health and disease is continually evolving, and the complexities of ligand binding and receptor activation warrant further exploration, including investigations into the effects of ligands in specific cellular contexts.

In conclusion, our study significantly advances our comprehension of dopamine receptor pharmacology. By deciphering the molecular basis of ligand-receptor interactions, we provide a basis for the design of more effective and targeted drugs for central nervous system disorders. This research contributes to the broader field of G protein-coupled receptor pharmacology and holds promise for the development of novel therapeutics that can improve the quality of life for individuals affected by these disorders. It is our hope that this work will stimulate further research in this domain, fostering innovation in drug development and improved patient care.

## Acknowledgement

Research reported in this publication was supported by the National Institute of General Medical Sciences of the National Institutes of Health under Award Number R01GM140078 (W.L.), the Advancing a Healthier Wisconsin Endowment (AHW) (L.O. and W.L.), and the Medical College of Wisconsin (MCW) start-up funds (L.Z. and W.L.). The protein samples were supplied by the protein production facility at MCW.

